# Dual role of Spreading Depolarization in the epileptic focus

**DOI:** 10.64898/2025.12.03.692116

**Authors:** Daria Vinokurova, Karina Tukhvatullina, Roustem Khazipov, Azat Nasretdinov

**Author notes:** **Correspondence:** Roustem Khazipov.

## Abstract

Spreading Depolarizations (SDs) are often associated with epileptic discharges. While SDs are traditionally thought contributing to the postictal depression and termination of epileptic discharges, seizures may also occur during SDs or may even follow SDs suggesting that interactions between SD and seizures are more complex. Here, we examined the interactions between SD and epileptic activity by spatially separating the epileptic focus and the site of SD initiation. Epileptic focus was induced by local intracortical injection of the potassium channel blocker 4-AP combined with the GABA(A) receptor antagonist gabazine, whereas extrinsic SDs were evoked by distal high potassium application. We found that extrinsic SDs promoted seizure-like events (SLEs) when the SD wave approached the epileptic focus, followed by suppression of epileptic activity when SD spread through the focus. The timing of SLE relatively SD varied at different recordings sites, with SLEs occurring before, during or after SD arrival depending on electrode position along the trajectory of SD propagation between the SD initiation site and the epileptic focus. During intracortical recordings, the proconvulsive effects of SD were associated with a wave of pre-SD neuronal excitation reaching the epileptic focus. The epileptic focus per se also demonstrated a resistance to the SD invasion. Thus, the interactions between SD and epileptic focus are not limited to postictal depression, and SDs may also promote epileptic activity in the hyperexcitable cortex.

**Key points:**

- Effects of SD on epileptic focus are dual: both pro– and anticonvulsive
- SDs promote epileptic discharges upon approaching the epileptic focus
- The timing of epileptic discharges relatively SD varies along the SD trajectory
- Proconvulsive effects of SD align with the pre-SD excitation
- Epileptic focus resists the SD propagation

## Introduction

Spreading depolarizations (SDs) are waves of collective depolarization of neurons and glial cells that spread slowly (at a rate of several millimeters per minute) throughout the gray matter of the cerebral cortex ^1–5^. Historically, SDs were first detected as waves of spreading depression of cortical electrical activity during studies of focal epileptic discharges induced by electrical stimulation of the cortex ^6^. Since then, a convincing body of evidence has been accumulated indicating a close association between SDs and epilepsy, both in patients ^7–16^ as well as in the experimental animal models in vivo ^8,9,11,17–25^ and in vitro ^26–31^ (for reviews, ^32–34^). The prevailing hypothesis is that SDs occurring during epileptic discharges lead to cortical inactivation as a result of depolarization block of neurons and cessation of epileptic activity, and SDs may thus be one instrumental in postictal depression ^18,19,22,35,36^. Besides, SDs are involved in seizure-induced cardiovascular arrest when it propagates to the brainstem causing SUDEP ^24^, cause headache in patients with epilepsy ^37,38^, underlie the antidepressant effects of electroconvulsive therapy ^8^, and cause postictal wandering in patients with temporal lobe epilepsy ^9^. However, in addition to the occurrence at the end of epileptic discharges and a clear contribution to their termination, SDs can also occur during the epileptic discharge, as in the case of partial superficial SDs ^17,39^ (see also ^40^), as well as before epileptic discharges as in the case of “spreading convulsions”, suggesting that SD may also promote epileptic activity ^6,7,10,13,29,31,41–46^. The wave of pre-SD excitation, which is associated with a moderate neuronal depolarization and neuronal excitation is a potential candidate for the proconvulsive effects of SD ^47–49^. On the other hand, epileptic cortex displays resistance to SD and SDs may even avoid penetration to the epileptic focus ^50,51^. Taken together, these observations suggest that the interactions between SD and paroxysmal activity are not limited to postictal depression, but are more sophisticated. However, the problem of interactions between paroxysmal activity and SD is complicated by the intricate spatiotemporal organization of both epileptic activity and SD initiation and propagation patterns. To circumvent these problems, we used multielectrode ECoG arrays and intracortical silicon probes to examine the interactions between SD and epileptic activity in a model, in which SD initiation site and epileptic focus were separated in space. We report dualism in the interactions between SD and the epileptic activity by showing that extrinsic SDs promote seizure-like events when SD approaches the epileptic focus, but terminate epileptiform discharges when SD fully invades the epileptic focus.

## MATERIALS AND METHODS

### Animal preparation

Wistar rats of both sexes aged from 3 to 8 weeks were used. Animals were prepared under isoflurane anesthesia at the surgical level (4% for induction, 2% for maintenance, Aerrane (Baxter, UK)). The skin and periosteum above both hemispheres and cerebellum were removed. Next, the neck muscles were detached from the occipital bone, then the wound was treated with bupivacaine (0.25%). The skull surface was dried and covered with cyanoacrylate glue. A metal bar was attached to the skull along the sagittal suture using cyanoacrylate glue and dental cement (Meliodent, Heraeus Kulzer, Germany). At the same time, isoflurane anesthesia was gradually reduced, and the animal was administered urethane (Sigma, USA) at the surgical level of anesthesia (1.5 g/kg, i.p.), confirmed by immobility, the absence of vocalizations and negative toe-pinch reflexes during the entire experiment. The animal was placed on a heated platform (37 °C, TC-344B; Warner Instruments, Hamden, CT) and the metal bar was attached to a fixation system via a ball joint. A craniotomy of 4×5 mm size centered on the left barrel cortex was performed with the detached bone piece left on site. To prevent herniation, the fourth ventricle was then punctured with a sharp needle. Then, a paper towel placed in a proximity to the puncture site was used to drain cerebrospinal fluid and lower intracranial pressure. Following this step, the bone piece was removed, and the dura mater was gently dissected along craniotomy perimeter with microscissors, then removed as a single flap. ECoG recordings of the local field potential (LFP) from the cortical surface were performed using custom ECoG electrode arrays (6 × 10 chromium-gold electrode grids on a polyimide film base with 0.4 mm separation distance between the electrodes and pre-fabricated holes of 0.2 mm diameter ^52^) placed epipially on the parietal cortex and covered with a liquid light paraffin (Panreac AppliChem, Spain) to reduce pulsations and to prevent cortex from drying. Intracortical recordings of local field potentials (LFP) and multiple unit activity (MUA) were performed using two linear multichannel silicon probes with iridium electrodes: 413μm^2^ surface area, 100μm separation distance (А1х16-5-100-413-А16, Neuronexus Technologies, USA). The probes were inserted vertically into the cortex through the holes in ECoG film to a depth of 1.6-1.8mm as follows: one probe was placed close to the site of 4-AP/gabazine injection whereas another one was placed more caudally at a distance of 1.6-2.0 mm from the first probe. The signals were amplified, lowpass filtered at 9kHz and digitized at 32kHz using a DigitalLynx SX amplifier and Cheetah 6.3.2 (Neuralynx, USA). Recordings were performed using full-band recordings with inverse filtering for signal reconstruction based on hybrid AC/DC-divider filter ^53^. A second cranial window ∼0.3mm in diameter for SD induction with epidural KCl (0.5M) application was made over the occipital cortex (∼6–7mm caudal and ∼5-6mm lateral from bregma). The KCl application chamber was built with a 1-2mm high dental cement wall around the cranial window ^54^. A cocktail of 4-aminopyridine (300 μM) and gabazine (30 μM) solved in PBS was applied using glass pipette (tip diameter 2–3 μm at 1 µl/min) connected to the 705 Hamilton syringe (50 μl, Hamilton, USA) filled with liquid light paraffin (Panreac AppliChem, Spain). The single injection volume did not exceed 0.5 µl and was carried using Micro4 Microsyringe Pump Controller (World Precision Instruments, USA) at cortical depth of 1000 μm.

### Data analysis

Raw data were preprocessed using a custom-developed suite of programs in the Matlab environment. Positive polarity is graphed as up throughout all manuscript. The original DC signal was downsampled to 1kHz and then used to characterize the LFP signal. SD onset was determined from the peak of the first derivative (SD’) of the 1Hz lowpass filtered LFP signal during the initial SD depolarization phase ^54^. Events with an SD’ peak <1mV/s were discarded from the analysis. SD stop depth was determined from the depth of the deepest electrode which displayed SD’ peak ≥1mV/s. Spectral power was valued using direct multi-taper estimators (1 Hz bandwidth, 3 tapers, 2 s spectral window with 0.1 s step, Chronux toolbox in Matlab) for the LFP signal on focal ECoG in the range of 2-70Hz (AC-band) during intervals: [-50 - 20] s before the SLE onset; between SLE onset and offset; 10 s after the SLE offset.

To detect population spikes (PSs), the first derivative of the average LFP (>0.1 Hz) was calculated across all ECOG channels (hereafter referred to as Common ECoG LFP’). PSs were then detected as local negative events below a threshold of –100 μV/ms with a minimum next event detection time of 40 ms. The frequency of detected PSs was calculated with a 1 s sliding window advanced in 1 ms steps. PS bursts with the PS frequency exceeding the 3 std threshold for more than 4 seconds were considered as SLE. The SLE onset and offset were defined as time of the PSs closest to the 1 std PS frequency threshold.

The borders of the epileptic focus were determined based on the ECoG LFP >0.1 Hz using the interpolated image of the PS amplitude (see Fig. 2A) as the points where the PS amplitude decreases to a level of 1/2 of the maximum value. The channel equidistant from the focus borders was taken as the center of focus along the x-projection of the ECoG matrix (10 channels). Then, based on the 2D coordinates of the focus borders, a circle approximation was performed, and the position of the focal center was refined. The obtained radius values were used to plot the average focus border and the corresponding probe positions.

For the extracellular multiple unit spike detection, the original wide-band signal was filtered (250-4000Hz, Daubechies wavelet filter ^55^), and negative local peaks >4 STD of the quietest 100s fragment in the control period (in the presence of 4-AP and gabazine before the KCl application) were considered as spikes. The MUA frequency was calculated in a sliding 100 ms window with a 1 ms step, whereas the total MUA frequency was calculated as the sum of MUA frequencies across all 16 channels of the probe. The location of the probe channel corresponding to the layer L5a was determined by the peak value of the vertical MUA frequency profile calculated for the period before the injection of 4-AP and gabazine.

To calculate the amplitude-temporal features of the slow SD wave, the original DC LFP signal was filtered in a range of 0.005-1 Hz, and for the resulting signal the peak negative value (SD amplitude) and the duration at half amplitude (SD half-width) were determined. The SD probability was calculated for each animal as the number of SDs observed at a given depth divided by the total number of SD episodes.

### Statistical analysis

Statistical analysis was performed using the Matlab Statistics toolbox. Wilcoxon rank sum and signed-rank two-sided tests were used to assess the significance of differences between samples. The level of significance was kept at p<0.05. Pooled data are presented as median, 25^th^ (Q1) and 75^th^ (Q3) percentiles, in the form of boxplots or violin plots.

### Study approval

The animal experiments were carried out in compliance with the ARRIVE guidelines. Animal care and procedures were in accordance with EU Directive 2010/63/ EU for animal experiments, and all animal-use protocols were approved by the Local Ethical Committee of Kazan Federal University (#24/ 22.09.2020).

## RESULTS

### Extrinsic SDs evoke SLEs in the epileptic focus

We explored the effects of SD on epileptic activity in cerebral cortex by separating in space the epileptic focus and the site of SD induction (Fig. 1A). Epileptic focus was formed in somatosensory cortex by intracortical injection of potassium channel blocker 4-AP and GABA(A) receptor blocker gabazine cocktail. Epileptic activity in the focus was characterized by regular interictal population spikes (PS) occurring at a frequency of 0.77 [0.61 1.00] Hz (median, [Q1 Q3]), and attaining 9.8 [6.1 10.0] mV (n=7 rats) on the ECoG electrodes close to the blockers injection site (Fig. 1B&C). SDs were evoked by focal epidural application of high-potassium (0.5-1 M) solution on visual cortex (VCx) at 3-4 mm from the epileptic focus. This induced recurrent SDs associated with characteristic negative DC shifts (Fig. 1B&C), and change in infrared light tissue transparency during concomitant IOS recordings. Remarkably, the vast majority of SDs (∼80%, Fig. 1D) were also associated with seizure-like events (SLE) manifested by bursts of PS with 15.4 [11.2 18.9] s duration (Fig. 1C, top panel), in which PS frequency rose to the peak value of 6 [4 7] PS/s (mean intraburst PS frequency of 1.9 [1.5 2.4]PS/s) (Fig. 1E). In some cases, SLEs were associated with a double-peak increase in PS frequency (Fig 1 B&C). SLEs were also associated with an increase in LFP power in the frequency range from 12 to 70 Hz (Fig. 1F). All SLEs were invariably followed by complete suppression of activity in the epileptic focus, characteristic of spreading depression causing postictal depression (Fig. 1B,C,E&F).

**Figure 1.**
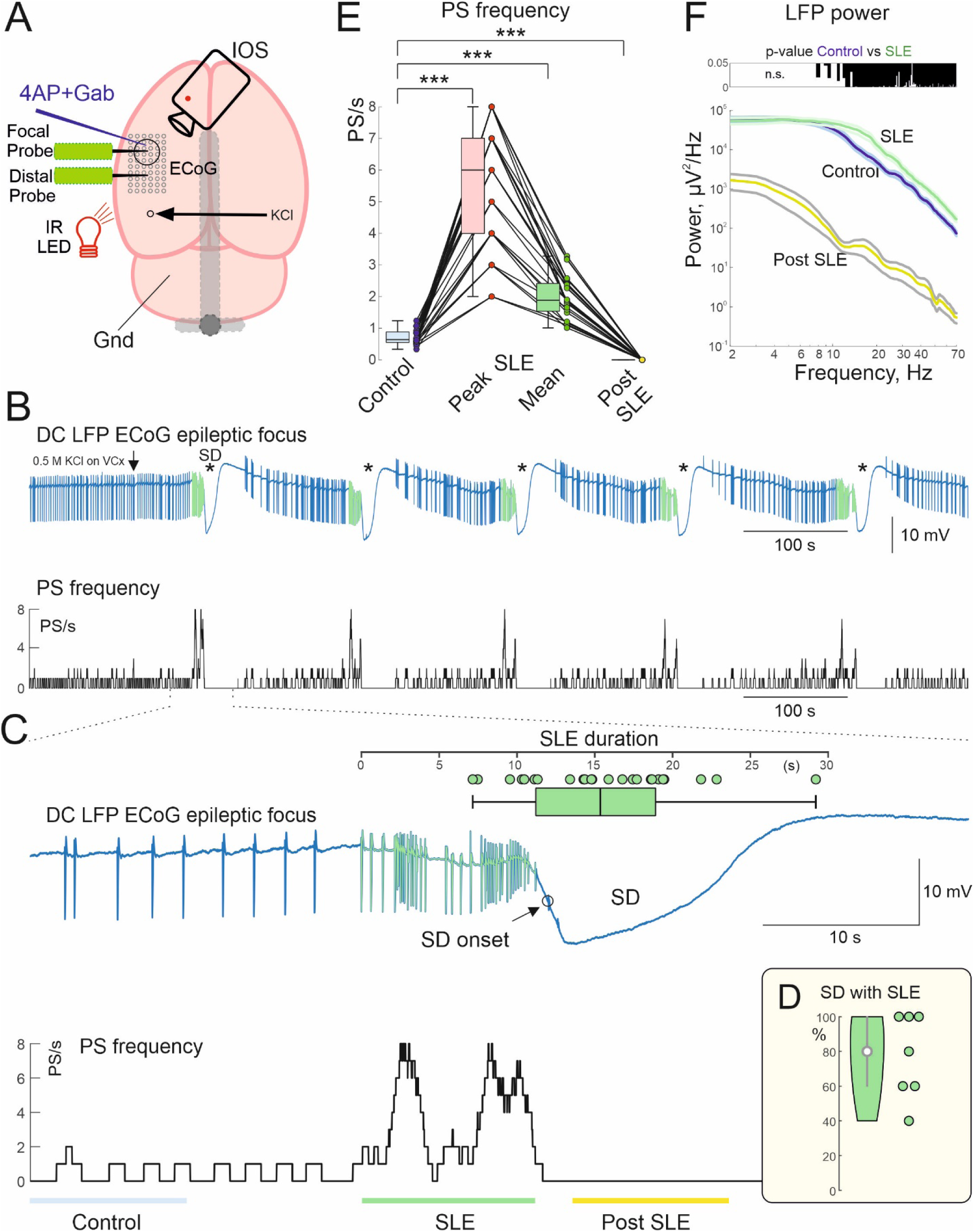
Extrinsic Spreading depolarizations (SDs) SD promote seizure-like activity in the epileptic focus. **(A)** Scheme of experimental setup for recordings of SDs evoked by distant epipial high-KCl application and activity in epileptic focus induced by local intracortical injection of the cocktail of 4-aminopyridine (4-AP) and gabazine. The gray bar represents the metal headfixation holder. **(B)** Cluster of five recurrent SDs (marked by asterisks) recorded from an epileptic focus. From top to bottom: DC-ECoG recordings from an electrode adjacent to the injection site (blue trace) and population spike (PS) frequency plot; seizure-like events (SLE) are colored green. (**C**) The first SD episode shown on expanded time scale. SLE duration (median with 25^th^ and 75^th^ percentiles) with time = 0 corresponds to the SLE onset is shown on top. Each dot corresponds to an individual SD. (**D**) Probability of SLE occurrence in association with SD. Each dot corresponds to an individual animal. (**E**) PS frequency before SD (Control, blue), the peak value during SD-associated SLE (pink), the mean value during SLE (green), and after SLE (yellow). (**F**) LFP power spectral density (on a logarithmic scale) in the 2-70 Hz range, calculated for the ECoG channel above the epileptic focus before SD (Control, blue), during SLE (green), and after SLE (yellow). Top: p – value plot for a difference between the LFP before and during SLE (twoway Wilcoxon signed rank test). **C-F**: Pooled data from n=24 SDs recorded from 7 rats.

**Figure 2.**
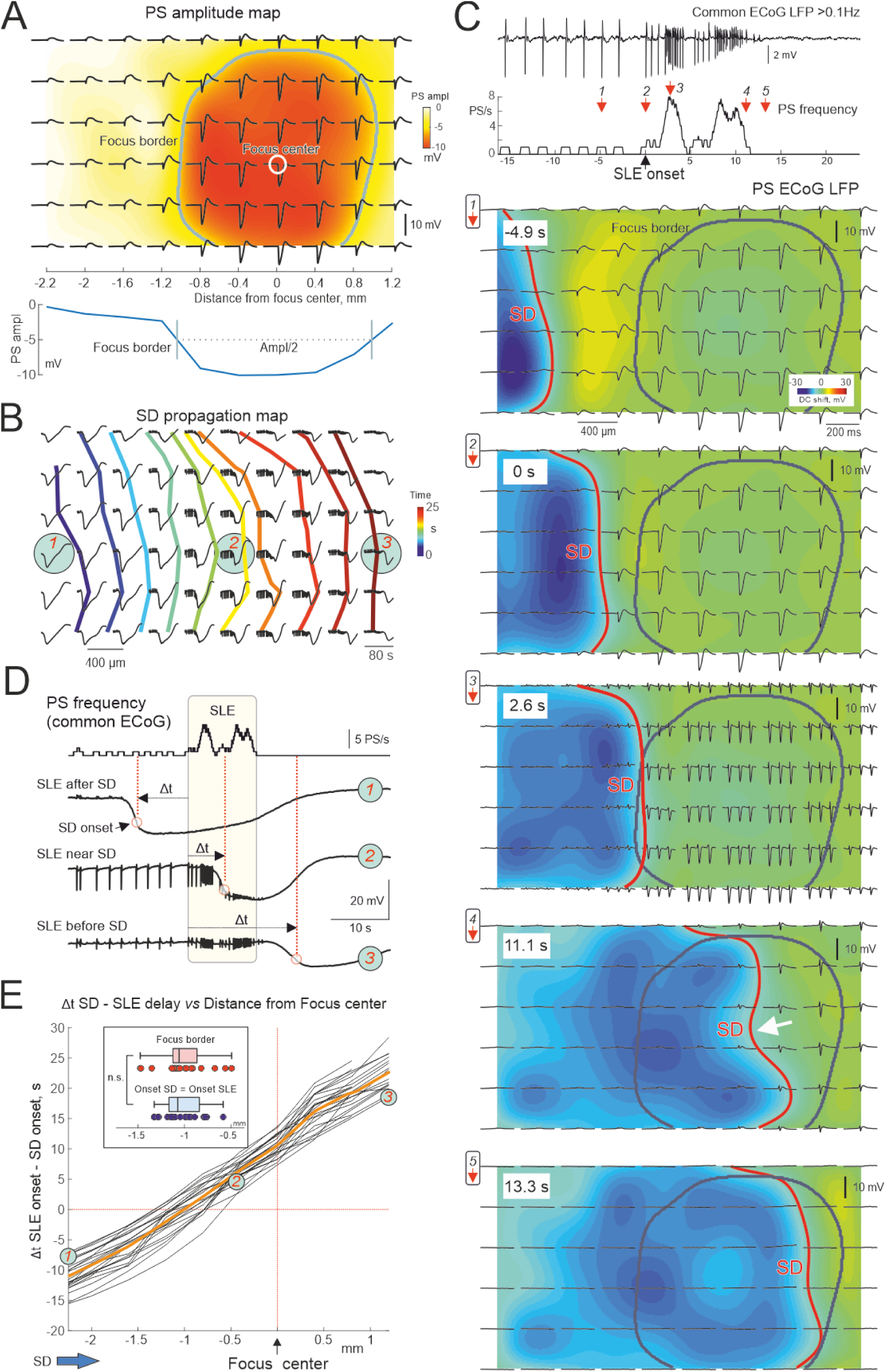
SD triggers SLE upon reaching the epileptic focus. **(A)** Average interictal PS on 60 ECoG channels (black) overlaid on the color-coded PS amplitude map. The center of the epileptic focus (white circle) is assigned to a channel with the maximal PS amplitude. The epileptic focus border (concentric grey line) delineates the region with > half-maximal PS amplitude as shown on the plot below. **(B)** DC-ECoG LFPs during SD induced by high KCl application 5 mm left from the array and propagating through the epileptic focus. Isochrones of SD onsets are indicated by color-coded lines. (**C**) Top, common ECoG LFP and corresponding PS frequency plot during high KCl-induced SLE. Below, examples of PSs (black) overlaid on color-coded DC-ECoG LFPs, and SD onset isochrones (red lines) at five time points indicated by red arrows on the PS frequency plot above. The focus border is indicated by the grey line. The time after distant KCl application is shown on the top left insets. Example #4: The white arrow shows the slowing of SD in the focus (**D**) PS frequency plot with DC-LFPs recorded from three selected ECoG electrodes on SD trajectory marked by cyan circles on panel B. Note that SD onsets (vertical dashed lines) occur before SLE (yellow rectangle) on the electrode 1, during SLE on the electrode 2, and after SLE on the electrode 3. (**E**) Relationship of the time lag of SD onset from the SLE onset (Δt) on the distance from the center of the epileptic focus in an ECoG row closest to the focus center. Each black line shows individual SD, the orange line shows the average. The highlighted circles 1, 2, and 3 correspond to the examples presented in panels B and D. The blue and red boxplots show the distance from the epileptic focus center where the SD and SLE onsets coincide (Δt = 0) and the distance from the focus borders to the focus center, respectively (p = 0.375, two-way Wilcoxon signed rank test), outliers are not shown **A-D**: Recordings from one rat, **E**: Pooled data from n=23 SDs recorded from 7 rats.

### SD triggers SLE upon approaching the epileptic focus

While the findings above indicate that extrinsic SDs promote SLE in the epileptic focus, they also raise a question of where is located the “hot-spot” of SD – triggered SLE. In this aim, we explored spatio-temporal dynamics of extrinsic SDs and SLE in horizontal cortical space. Using ECoG multielectrode arrays, we first determined spatial features of the epileptic focus relying on the amplitude of PS on different ECoG channels. As shown on the PS amplitude map on Fig. 2A, PS amplitude was maximal on about 5 channels around the site of the blockers injection and decreased at more remote channels. Epileptic focus border was defined at halfmaximal PS amplitude revealing round-shape epileptic focus of 0.94 [0.74 1.09] mm diameter with a center in the blockers injection site. These metrics of the epileptic focus were assessed prior to each SD. Importantly, PSs were highly synchronized on all electrodes of the ECoG array (Fig. 2A), and therefore common average ECoG signal was used for PS and SLE detection. Next, we assessed SD propagation by monitoring onset of SD-related DC shifts on ECoG electrodes and concomitant IOS recordings. As shown on a typical example presented on Fig. 2B, SD propagated from the left to the right relatively epileptic focus through the entire recording field. On average, SD propagated at a speed of 5.9 [5.4 6.3] mm/min, but some anisotropy in SD propagation was observed when SDs approached the epileptic focus (see below). Snapshots illustrating epileptic activity on ECoG array at different times of SD propagation through the ECoG array are shown on Fig. 2C (Time of SLE onset = 0). SD propagation is indicated by blue-coded negative DC shift, whereas SD front is indicated by the red line. Interictal activity remained unchanged while SD propagated through the cortex distant from the epileptic focus (Fig. 2C, T= –4.9 s), and SD in these recording sites largely preceded SLE (Fig. 2B,D, ECoG#1). However, as soon as SD attained the border of epileptic focus, it triggered SLE (Fig. 2C, T = 0 s and 2.6 s), with SD in these recording sites occurring close to the SLE onset (Fig. 2B,D, ECoG#2). Further SD propagation through the epileptic focus was associated with the suppression of SLE (Fig. 2C, T = 11.1 s and 13.3 s), and SD in these recording sites occurred after SLE (Fig. 2B,D, ECoG#3). We further quantified the delay between the SLE and SD onsets (Δt SD-SLE) as a function of a distance from the center of the epileptic focus along the SD trajectory towards the center of epileptic focus (row 4 on Fig. 2B,C; Fig. 2E). At the left electrodes (near the SD initiation site and distant from the epileptic focus), SLEs were strongly delayed after SD. At electrodes near the border of the epileptic focus, SLEs occurred with minimal delay after SD. In the center of the epileptic focus and at remote electrodes along the SD propagation path, SLEs largely preceded SDs (Fig. 2E). On average, SLE onset was observed at the time when SD attained a distance of –1.07 [-1.17 –0.84] mm from the center of the epileptic focus, matching the location of the epileptic focus border (–1.06 [-1.12 –0.87] mm) on the SD-entry side (Fig. 2E, top boxplots). The conclusion from these findings is two-fold: (i) the timing of SLE onset from SD largely varies among cortical sites depending on a distance from the epileptic focus, and (ii) SLE is initiated when SD approaches the epileptic focus and attains its border. Of note, epileptic activity was suppressed in the regions invaded by SD both in the epileptic focus and beyond, and following few minutes of recovery from SD, the interictal activity returned to pre-SD levels.

### Extrinsic SD terminates SD-induced SLE

While the SD-induced SLE onset was simultaneous on entire ECoG array, the termination of SLE and SLE duration varied between the electrodes. As shown on example recordings from a row of electrodes on SD trajectory from the SD initiation site to the epileptic focus (Fig. 3A), SLE had terminated much earlier and lasted for a shorter time on the electrodes located closer to the SD initiation site, whereas at the electrodes located closer to and within the epileptic focus, SLE terminated later and lasted for a longer time (Fig. 3A,B). Of note, short duration SLE at the electrodes located outside the epileptic focus proximal to the SD initiation site (ECoGs #2-4 on Fig. 3A) occurred after the SD entry to these locations that is reminiscent of “spreading convulsions” phenotype ^10^. Similarly, SLE also persisted after SD onset for up to ten seconds at sites located at the border and within the epileptic focus proximal to the SD entry (ECoGs #5-6 on Fig. 3A). Post-SD SLEs at these locations were likely supported by the activity in deep layers, which are invaded by SD at a delay from the superficial layers (see below). Increase in SLE duration along the SD propagation trajectory was evident on entire ECoG array (Fig. 3C). SLE termination occurred when SD invaded most of the epileptic focus and attained 0.35 [0.26 0.64] mm from the focus center along the SD trajectory, closer to the SD exit border from the epileptic focus (Fig, 3D, see also Fig. 2C, T=11.1 s and 13.3 s). Together, these findings suggest that SLE duration strongly varies along the SD trajectory and that SD both initiates SLE upon reaching the border of the epileptic focus and terminates SLE when invades most of the territory of the epileptic focus, and when the critical mass of the epileptic network required for SLE is suppressed by SD.

**Figure 3.**
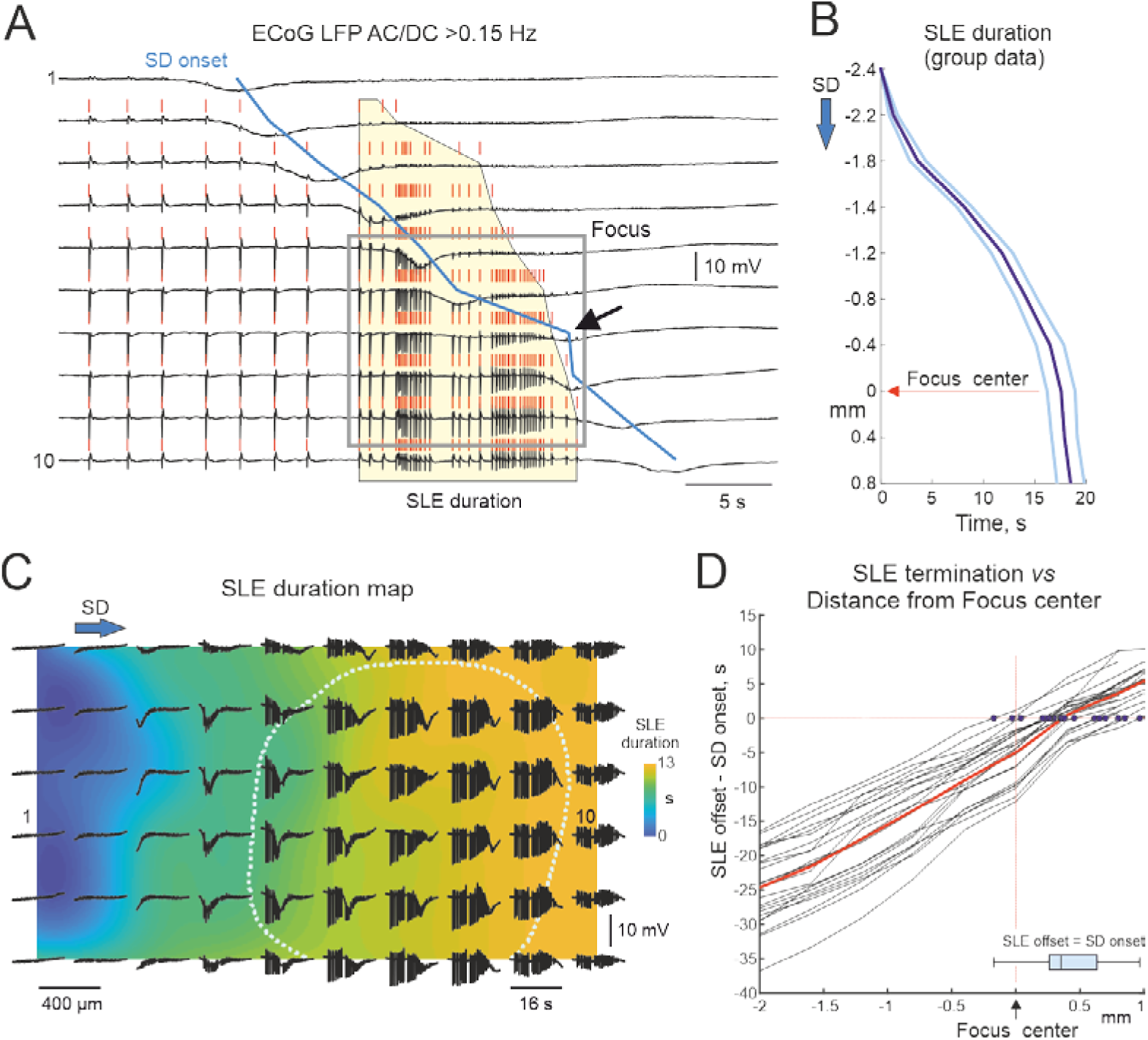
SD terminates SLE and determines SLE duration along the SD trajectory. **(A)** Example recordings of AC-ECoG LFP (black) in a selected electrode row on a SD trajectory from SD initiation site to the center of the epileptic focus, PSs (vertical red tics), SD onsets (blue line), focus border (grey box). Note that SLE (yellow figure) terminates earlier and SLEs have shorter duration at the electrodes closer to the SD initiation point (top channels). The black arrow shows the slowing of SD in the focus. (**B**) Dependence of SLE duration on the distance from the focus center along the trajectory of SD propagation (mean ± standard error). (**C**) Examples of AC-ECoG LFP during SD-induced SLE overlaid on the color coded SLE duration map, border of the epileptic focus is outlined by dashed grey line. (**D**) The dependence of the time lag of SLE termination from the SD onset on the distance from the epileptic focus center. The blue boxplot shows the distance from the epileptic focus center where the SD onset coincides with the SLE termination, that is, the SD coordinate where SLE terminated. A, C: Recordings from one rat. B, D: Pooled data from n=24 SDs recorded from 7 rats.

### SD and SLE through the cortical depth

Since SD spreads not only in horizontal, but also in vertical cortical space ^17,49,54,56–59^, we further explored the relationships between SD and SLE by recording activity through the entire cortical depth using two linear 16 channel silicon probes. One of them was inserted into the “naïve” cortex on a way of SD towards epileptic focus (“distal” probe), while another probe was inserted into the epileptic focus (“focal” probe, Fig. 4A&B) with concomitant ECoG recordings. In agreement with the ECoG data (Fig. 2A), the amplitude of interictal PSs was maximal at the focal probe, with the largest PS amplitude observed in the superficial layers (Fig. 4C). Extrinsic SDs sequentially occurred first on the distal probe and then on the focal one, with the SD – DC shifts on the top probes’ channels similar to the SD – DC shifts recorded from the nearby ECoG electrodes (Fig. 4D,E). In keeping with the top-down gradient of vertical SD propagation ^17,49,54,58,59^, SDs spread from the superficial to deep layers on both probes (Fig. 4D). As a result of the vertical SD propagation, the timing of SLE relatively SD also varied depending on cortical depth. As shown in the example recordings presented on Fig. 4D, SD started earlier than SLE on the top channels of the distal probe, whereas SD and SLE onsets coincided in time at the middle of cortical depth of the distal probe. On the other hand, SD arrived about ten seconds after the SLE onset according the data from the deepest channels (Fig 4D&E). Therefore, SD started after SLE onset at all cortical depths at the probe in the epileptic focus, but the SD-SLE delay also strongly increased with cortical depth, so SD started before and after SLE termination in the superficial and deep cortical layers, respectively (Fig 4D&E). The diversity in SD delays from the SLE onset and their increase with cortical depth was evident at the group data level (Fig. 4F).

**Figure 4.**
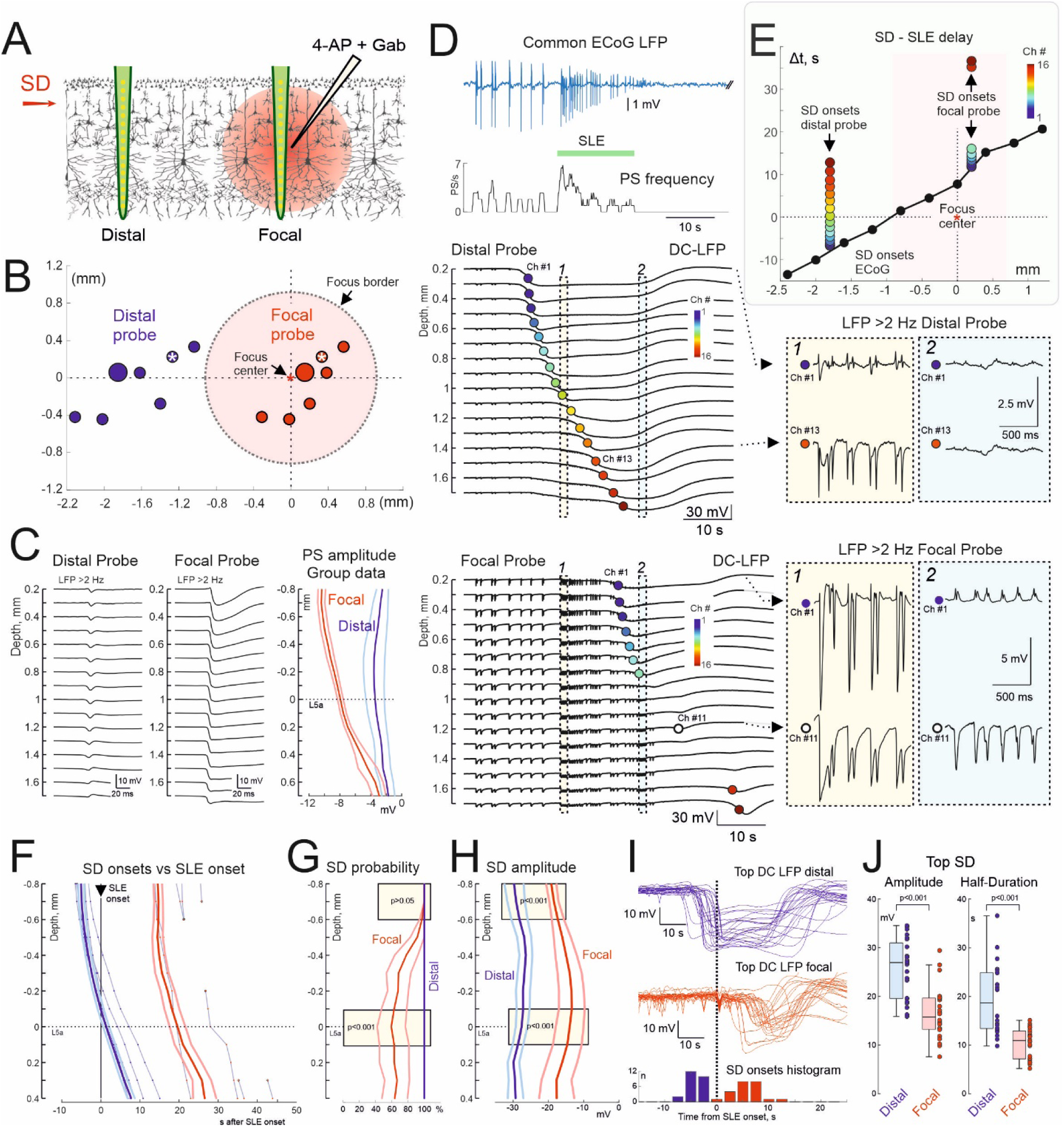
Delay of SLE from SD depends on cortical depth. **(A)** Sideview scheme of dual DC-LFP recordings across cortical depth using one silicon probe inserted distally from the epileptic focus closer to the SD induction site, and the second probe inserted into the epileptic focus. (**B**) View from above of the distal (blue) and focal (red) probes’ location relative to the epileptic focus center for all experiments, gray circle represents the grand average epileptic focus border (n=7 animals). (**C**) Depth profile of a PS at distal (left) and focal (right) 16 channel probes. On the right, PS amplitude at different cortical depths (mean ± standard error) at distal (blue) and focal (red) probes. (**D**) Top-down: Left, Common AC-ECoG LFP and PS frequency plot, and DC-LFP recordings through the cortical depth on the distal (middle) and focal (bottom) probes. Colored circles indicate the SD onsets, color codes the channel number. Right, examples of AC-LFP recordings from the channels #1 and #13 the distal probe and channels #1 and #11 of the focal probe during the two highlighted time segments (Episodes 1 and 2) on expanded time scale. (**E**) Dependence of the time lag of SD onset from the SLE onset on the distance from the center of the epileptic focus in an ECoG row closest to the focus center (black) and at the color-coded cortical depth of the distal and focal probes shown on the panel D. (**F**) Depth profile of SD onset at distal (blue) and focal (red) probes. Four examples are shown for one animal, group data are presented relative to the channel in layer L5. Open circles indicate the SDs interrupted in the middle of the cortex. (**G**) Depth profile of SD occurrence across cortical depth. (**H**) Depth profile of the SD amplitude in focal (red) and distal (blue) probes (mean ± standard error). (**I**) Superimposed DC LFPs from the top channel of the distal (blue) and focal (red) silicon probes and corresponding histogram of SD onsets relative to SLE onset. (**J**) Amplitude and duration of SDs at the top channel of the distal and focal probes (p=7.5e-4 and p=4.9e-5, respectively; twoway Wilcoxon signed rank test). A: The cortex texture is adapted from the Ref ^64^. E-J: Pooled data from n=24 SDs recorded from 7 rats.

During their vertical propagation, SDs created transient states with unique organization of epileptic activity within a cortical column. Two examples of such transient states, during which SD attained the middle depth at the distal probe (Example 1) and the focal probe (Example 2), respectively, are shown on Fig. 4D (right panels). On example 1, PSs were suppressed or even changed the polarity from negative to positive in the superficial layers already invaded by SD (Ch#1) while maintained negativity in deeper layers not yet invaded by SD (Ch#13) on the distal probe, as well as at all depth of the focal probe (for comparison with the pre-SD state, see Fig. 4C). On example 2, epileptic activity was blocked by full SD across all channels of the distal probe, whereas on the focal probe, PSs switched the polarity from negative to positive in the superficial layers invaded by SD (Ch#1) while maintaining negativity in deeper layers (Ch#11). The changes in the electrographic phenotype of the PSs during vertical SD propagation were also associated with a suppression of neuronal activity (MUA) in the superficial but not deep layers (Fig. 5A,B).

**Fig. 5.**
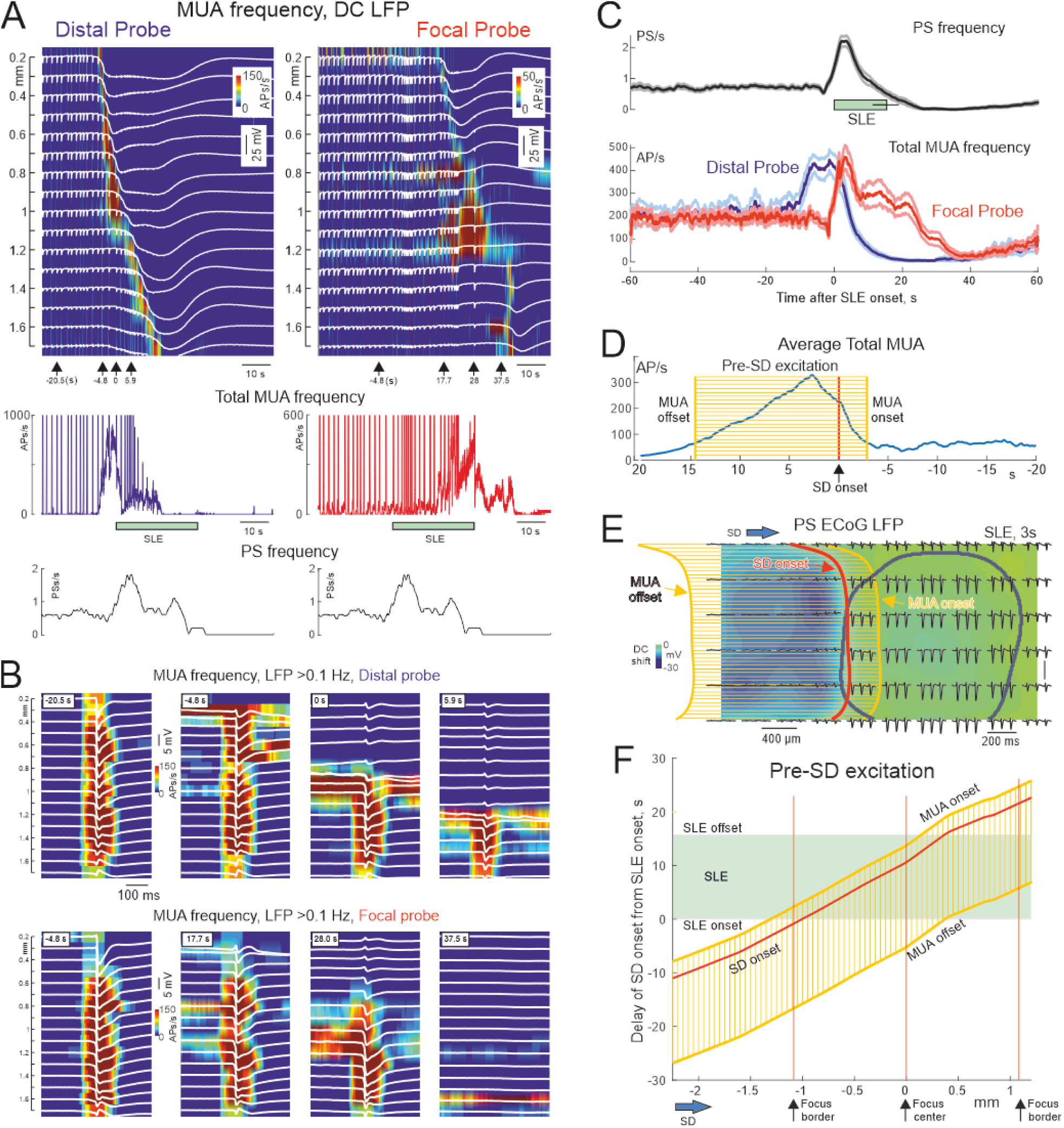
Wave of pre-SD excitation and SLE. **(A)** Color-coded MUA density map across cortical depth during SD recorded by distal (left) and focal (right) silicon probes. White traces – DC-LFP at different cortical depth. Silicon probes position is indicated by white asterisks on Fig. 4B. Below, corresponding total MUA frequency across all depth and common ECoG PS frequency plots. (**B**) Example PSs at different time points from SLE onset (as indicated by bottom arrows on panel A) along SD propagation through the cortical column recorded by distal (top) and focal (bottom) probes. (**C**) Grand average of common ECoG PS frequency (top) and total MUA frequency (bottom) at distal (blue) and focal (red) probes aligned to the SLE onset. (**D**) Grand average of total MUA frequency aligned to the surface SD onset. Boxplot shows the pre-SD total MUA burst onset and offset times. (**E**) Reconstruction of the pre-SD excitation (yellow grid) at the 3^rd^ second of SLE in the horizontal cortical plane. Template of pre-SD excitation (panel D) was triggered by surface SD onset during ECoG recordings (SD onset, red line). DC-ECoG LFP is coded by background color (blue, SD-related negative DC shift). Black traces – 200 ms fragments of ACECoG LFP. Grey line – epileptic focus border. (**F**) Propagation of the reconstructed pre-SD excitation wave in coordinates of SD – SLE onsets delay (Y-scale, grey area shows the entire SLE period) along the SD trajectory from the SD initiation site through the center of the epileptic focus (X-scale, focus center and borders are indicated by bottom arrows and vertical red lines). C-F: Pooled data from n=24 SDs recorded from 7 rats.

Interestingly, while full SDs on the distal probe displayed vertical propagation profile with SDs sequentially invading all layers similar to that in normal cortex, SDs in the epileptic focus showed a tendency to avoid middle cortical depth with eventual interruption of SDs in the middle layers near the depth of the proconvulsants injection (Fig. 4D, F). During group analysis vertical position of the silicon probe channels was normalized to the depth of the layer 5, which hallmark is the highest neuronal firing level ^60–63^. Tendency of SD avoiding middle layers of the focus manifested in lower SD probability in the middle and deep layers compared to the superficial depth in the focus and to the distal probe (Fig. 4F,G) and the smallest SD amplitude at the middle depth of the focus (Fig. 4H). Similar tendency of SD to avoid the epileptic focus was also supported by smaller SD amplitude through the entire depth of the focus compared to distal site (Fig. 4H), and smaller SD amplitude and shorter SD duration at the top channels of the silicon probes in the epileptic focus (Fig. 4I,J). Moreover, a slowing of SD at the focus was also observed during propagation in the horizontal plane (Fig. 2C Example #4 and Fig. 3A, arrows). While certain degree of damage to the cortex by injection of blockers cannot be excluded, depth profile of PS in the focal probe (Fig. 4C) as well as persistence of MUA in middle cortical layers in the core of the focus (Fig. 5A,B) favor the idea that epileptic cortex shows resistance to SD that is in agreement with findings in other epilepsy models and in the human epileptic cortex^50,51^.

### Pre-SD excitation and SLE

The present findings indicate that the initiation of SLE is associated with SD entering the epileptic focus. What are the mechanisms underlying seizures triggered by SD in the epileptic focus? While SD causes nearly complete loss of membrane potential, depolarization block and full suppression of all forms of activity, these phenomena do not occur not in an all-or-none manner but gradually. Indeed, SD is preceded by a pre-SD excitation phase, characterized by moderate, near-threshold neuronal depolarization and increased neuronal activity ^19,47,49,54,56,65^, so pre-SD excitation could be potentially involved in the proconvulsive effect of SD in the epileptic focus. Therefore, we further explored spatial-temporal organization of neuronal activity during spread of SD through the epileptic focus using extracellular silicon probe recordings of multiple unit activity (MUA) (Fig. 5). We found that SDs are preceded by a wave of pre-SD excitation firstly on the distal and then on focal silicon probes, characterized top-down temporal organization corresponding to the vertical SD propagation (Fig. 5A, B). While neuronal firing during pre-SD period was almost completely synchronized in PSs, pre-SD excitation was also associated with elevation of MUA in the inter-PS intervals, along with MUA suppression during both PSs and inter-PS intervals in the layers invaded by SD (Fig. 5B). While the pre-SD excitation, assessed as total MUA across all layers of the cortical column, largely overlapped in time with SLE, it was delayed in the focal probe from the distal probe along with a horizontal SD propagation (Fig. 5C). We further attempted to characterize the pre-SD excitation dynamics during SD propagation through the epileptic focus in horizontal cortical plane. In this aim, we built a template of pre-SD excitation based on total MUA assessment using the silicone probe recordings merging data from the distal and focal probes relatively to SD onset detected on the cortical surface (Fig. 5D). In the horizontal plane, although pre-SD excitation within a column (total duration 17.4 s) started shortly before the surface SD (–2.9 s), the peak and vast majority of pre-SD excitation occurred after the SD onset at the cortical surface (+14.5 s, Fig. 5D). We next plotted the template of pre-SD excitation on the map of horizontal SD propagation taking SD onsets detected on ECoG electrodes as local time reference points (Fig. 5E). This experimental setup enabled to reconstruct propagation of the pre-SD excitation wave in horizontal cortical plane (a reconstruction at the third second of SLE is shown on Fig. 5E), and to relate pre-SD excitation to the coordinates of the epileptic focus in horizontal cortical space, as well as to the SLE onset and offset in time. Results of such analysis in a reduced dimension of SD trajectory from the SD initiation site through the center of the epileptic focus are shown on Fig. 5F. It is noticeable that the SLE onset is associated with entry of pre-SD excitation wave to the epileptic focus, and that propagation of the pre-SD excitation through the epileptic focus occurs through the entire time course of SLE. It may seem paradoxical that pre-SD excitation within the focus culminates at the end of SLE. However, pre-SD excitation wave is paralleled by a wave of SD-related suppression of activity, which propagates from superficial to deep layers (Fig. 5B,C), reducing the mass of the epileptic network. Therefore, it is plausible that SLE termination is caused by SD-induced depression of activity within a critical mass of the epileptic network despite of persisting pre-SD excitation in the deep layers.

## DISCUSSION

Our main finding is that SD exerts a dual role in the epileptic focus: while extrinsic SDs promote epileptiform activity when the SD approaches the epileptic focus, they also terminate seizures when SD fully invades the epileptic focus. These findings also provide mechanistic explanation for the diversity in temporal relationships between SD and SLE, with SLE following SD, occurring during SD and preceding SD at different cortical sites (both in horizontal and vertical space) relatively the trajectory of SD propagation.

The present findings suggest the following model of the complex and dynamic interactions between SD and epileptic focus. Activity in the hyperexcitable network of epileptic focus is organized in regular interictal PSs at the resting state prior to SD. Wave of SD, generated far away from the focus slowly propagates toward the epileptic focus causing a sequence of pre-SD excitation followed by depression in the cortical territories recruited by SD ^47–49^. Furthermore, pre-SD excitation does not cause epileptic activity in the “naïve” cortex, where inhibition efficiently balances excitation and shapes neuronal network excitation in low-amplitude gamma oscillations ^27,49,65^. Of note, in the hippocampus the prodromal pre-SD excitation may be organized in brief and local large amplitude gamma PS bursts ^48,66,67^. At this stage, activity in the epileptic focus remains unchanged, and SD in these regions distant from the focus precede forthcoming SLE. However, when SD approaches the focus border, pre-SD excitation starts interacting with the hyperexcitable focus network causing an increase in the PS frequency resulting in SLE. The mechanism of pro-convulsive effect of SD is likely to be fundamentally similar to that of pre-SD excitation, and involves mild neuronal depolarization at the SD onset caused by elevation of extracellular potassium, increase in neuronal firing and activation of local excitatory connections and non-synaptic glutamate release, as well as ephaptic interactions ^1–3,68,69^. Of note, the levels of overall increase in neuronal firing during pre-SD excitation phase in the naïve cortex and in the focus are similar. However, unlike relatively weak and local neuronal synchronization in gamma oscillations in the naïve cortex, pre-SD excitation causes SLE in the epileptic focus where inhibition is suppressed ^70–73^. The transition from low voltage fast activity at the SD onset to SLE is also reminiscent of similar paradigm at the seizure onset ^34,74–78^ raising a question of similarities in the generative mechanisms of low voltage fast oscillations at seizure and SD onset.

SLE lasts as long as pre-SD excitation propagates through the epileptic focus, followed by wave of spreading depression caused by neuronal depolarization above the action potential depolarization block resulting in SLE termination, as well as postictal depression of interictal PSs. SLE terminates when SD invades most of the epileptic focus suggesting that a critical mass of neuronal network in the focus is required for SLE generation. While SLE onset is highly synchronous in the entire epileptic focus, SLE termination is a dynamic process determined by SD propagation both in horizontal and vertical cortical space. Horizontal SD propagation through the epileptic focus creates transient states during which epileptic activity is suppressed across entire cortical depth while SLE persists in the regions of the epileptic focus in front of SD propagation. As a result, SLE duration differs within the focus, being shortest in the regions first invaded by SD and longest in more distant regions not yet invaded by SD along the SD trajectory. In keeping with the top-down gradient in SD propagation across cortical depth, SD first recruits superficial layers that also creates transient network states in vertical dimension. While PS normally involve the entire cortical column, SD first suppresses PSs in the superficial layers but epileptic activity persists in deep layers. Change in PS from negative to positive polarity is a hallmark of this transient state with the partial SD invasion to the superficial layers, that has been previously described in case of partial superficial SDs in the flurothyl mode of generalized seizures ^17^. This phenomenon likely involves a suppression of superficial generator, which predominantly contributes to PS negativity at the cortical surface, and unmasking of the positive passive source of the deep PS generator, that is mechanistically similar to a boost in surface delta power during partial superficial SDs ^49^.

We found that the core of the epileptic focus is resistant to SD propagation. This is in keeping with previous findings in other models of focal epilepsy including local picrotoxin, in which extrinsic SDs similarly avoided penetration to the epileptic focus ^51^. Similarly, slices of chronically epileptic human and rat neocortex display a resistance against SD in vitro ^50^. Interestingly, resistance to SD was manifested not only by a reduced SD amplitude and duration, as well as by slowing down of SD propagation in the epileptic focus at the cortical surface, but also by low penetration rate and smaller SD amplitude in the core of the epileptic focus, at around the depth of intracortical injection of the proconvulsants cocktail. In some cases, we observed quite unusual “sandwich” – like SD phenotype with SDs occurring only in the superficial and deep layers but avoiding middle cortical depth of the epileptic focus, that is the very core of the epileptic focus. While the mechanisms of higher resistance of the epileptic cortex to SD remain to be elucidated, resistance of the epileptic focus to SD should be considered during SD detection in the epileptic patients.

Thus, the results of the present study suggest dynamic and complex interactions between SDs and the epileptic focus, in which SDs exert dual pro– and anti-convulsive roles. While SDs clearly contribute to the postictal depression and termination of epileptic discharges, they may also promote seizures in the epileptic focus through the synergistic interaction of pre-SD excitation with hyperexcitable networks of the epileptic focus. Our results also point to highly dynamic nature of the interactions between SD and epileptic focus, both in horizontal and vertical cortical space, that creates unique transient network states with distinct electrophysiological phenotypes. Present assessments of the network function during the interactions between SD and epileptic focus, and their electrophysiological manifestations in three-dimensional cortical space, could be useful in clinical investigations and interpretation of the electrophysiological data in epileptic patients and in other pathological states associating SD and epileptiform activities including brain trauma, ischemia, tumor and migraine.

## DATA AVAILABILITY

Animal data supporting the results of the study are available from the authors upon request.

## CODE AVAILABILITY

Codes for the data analysis are available from the authors upon request.

## ACKNOWLEDGEMENTS

The research was supported by the RSF grant 22-15-00236-P

## Notes

### Competing Interest Statement

The authors have declared no competing interest.

